# Acetyl-CoA Synthetase Mutations S868G and V949I Do Not Confer Resistance to Antimalarial Drugs *in vitro* in *Plasmodium falciparum*

**DOI:** 10.1101/2024.06.03.597226

**Authors:** Wei Zhao, Zheng Xiang, Weilin Zeng, Yucheng Qin, Maohua Pan, Yanrui Wu, Mengxi Duan, Ye Mou, Tao Liang, Yanmei Zhang, Cheng Liu, Xiuya Tang, Yaming Huang, Gongchao Yang, Liwang Cui, Zhaoqing Yang

## Abstract

*Plasmodium falciparum* acetyl-CoA synthetase (PfACAS) protein is an important source of acetyl-CoA. We detected the mutations S868G and V949I in PfACAS by whole-genome sequencing analysis in some recrudescent parasites after antimalarial treatment with artesunate and dihydroartemisinin-piperaquine, suggesting that they may confer drug resistance. Using CRISPR/Cas9 technology, we engineered parasite lines carrying the PfACAS S868G and V949I mutations in two genetic backgrounds and evaluated their susceptibility to antimalarial drugs in vitro. The results demonstrated that PfACAS S868G and V949I mutations alone or in combination were not enough to provide resistance to antimalarial drugs.

## INTRODUCTION

Malaria, an infectious disease caused by *Plasmodium* parasites and transmitted by mosquitoes, remains one of the most significant global public health problems. *P. falciparum* has shown reduced sensitivity to artemisinin-derived antimalarials in Southeast Asia^1-8^ and Africa^9-15^. Clinical artemisinin resistance (ARTR) has been linked to mutations in the propeller domain of the *pfk13* gene^16-17^. However, despite this well-established connection, the current molecular marker for ARTR, the *pfk13* gene, appears to provide an incomplete explanation for the complex problem of ARTR. Some evidences showed that the parasites exhibited ARTR even in the absence of mutations in the *pfk13* gene, which suggests that factors beyond the *pfk13* gene may also play a role in conferring resistance to artemisinin in *P. falciparum*^18-19^. Therefore, it is urgent to explore and identify new molecular markers associated with antimalarial drug resistance.

In our previous study, we collected 11 cases of *P. falciparum* recrudescence from Chinese migrant workers who returned from African regions^20^. These Chinese migrant workers are a unique group, who had resided in Africa for an extended period and encountered multiple episodes of malaria, despite malaria prophylaxis by self-administering artemisinin either at home or treatment at local clinics. These individuals experienced a recurrence of malaria symptoms within 28 days after treatment with intravenous artesunate (AS) and orally dihydroartemisinin-piperaquine (DHA-PPQ), the treatment mode usually used for severe malaria^21^, during their hospitalization at Shanglin Hospital, China^22^.

We compared 11 recrudescent *P. falciparum* with 13 concurrent cured cases and 110 cases of African origin downloaded from SRA database (Sequence Read Archive, https://www.ncbi.nlm.nih.gov/sra) by whole genome sequencing analysis, and found mutations in PfACAS S868G and V949I in two recrudescent strains, accounting for 18% (2/11). PfACAS gene encoding ACAS enzymes that catalyze the condensation of acetate and CoA into acetyl-CoA is a key enzyme in many organisms and acts at the central point of energy metabolism such as Kreb’s cycle, lipid, and phospholipid synthesis^23^. In the plasmodial genome, only one PfACAS protein has been identified and it is an important source of acetyl-CoA. Prata^24^ found that knockout or inhibition of the PfACAS gene had an impact on the growth of malaria parasites, PfACAS serves as a vital cellular metabolite that is indispensable for the survival of these parasites. Summers^25^ described that several mutations in the PfACAS gene of *P. falciparum* (A597V, T648M, Y607C, and A652S) conferred resistance to two small molecules of antimalarial compounds (MMV019721 and MMV084978). In our study, we found mutations S868G and V949I in PfACAS by whole-genome sequencing analysis in some recrudescent parasites after antimalarial treatment. We suspected that these two mutations may confer resistance to these African parasites. Unfortunately, mutations S868G and V949I in PfACAS of African *P. falciparum* strains were not adopted in our laboratory, thus rendering the resistance phenotype unattainable in our research. To verify this view, we engineered parasite lines carrying the PfACAS S868G and V949I mutations in two genetic backgrounds, including 3D7 and the similar background strain 16-129 (from the special group, but no recrudescent strains were found, had visited the same local place in Africa and might got infection at similar time), and evaluated their *in vitro* susceptibility to antimalarial drugs.

## MATERIAL AND METHODS

### Plasmid construction and preparation

Based on pL6CS-Xu-HDHFR1 and pUF1-Cas9-BSD plasmids, we constructed pL6CS-Xu-gRNA-Control/S868G/V949I/S868G+V949I plasmids, which offer donor DNAs and gRNA. The donor carrying S868G or V949I was amplified using primers listed in Table S2. All primers bore the desired mutations or 20bp necessary for homologous arms. All PCR amplifications were done with high-fidelity polymerase (Vazyme) following the recommended protocols. Subsequently, the pL6CS-Xu-gRNA plasmid and donor were digested by restriction enzymes *Xhol I* and *Avr II*. The products were purified, and the linearized plasmids and donor fragments were mixed in a recombinant enzyme system (Vazyme) to construct pL6CS-Xu-gRNA-Control/S868G/V949I/S868G+V949I plasmids, respectively. After that, the plasmids were transformed into XL-10 gold competent cells respectively. Plasmids were extracted using Plasmid Mini Kit (Omega) and checked with sequencing to obtain correct plasmids. Additionally, we also fused a 3×HA tag to the strains at the terminus of PfACAS. Primers required for this construction are provided in Table S3.

### *P. falciparum* culture and transfection

Parasites were cultured in type O erythrocytes and grown with complete medium containing 10.4g/L RPMI 1640, 2g/L NaHCO3, 0.1 mM hypoxanthine, 50mg/L gentamycin and 5g/L Albumax II, 25mM HEPES, and were incubated at 37°C in a gas mixture of 5% CO_2_, and 5% O_2_^26^. The pL6CS-Xu-g-ACAS and pUF1-Cas9-BSD plasmids construction was carried out according to the previous report^27^. Transfection was performed by pre-loaded^28-29^. Briefly, Parasites were synchronized using 60% Percoll gradients to capture schizonts for transfection, and fresh red blood was washed with 1× cytomix solution just before transfection. The transfection mixture contained 100 μg pUF1-BSD-Cas9 plasmid, and 100 μg pL6CS-Xu-g-ACAS plasmid, 2× cytomix (the volume is consistent with the plasmid volume), 160 μl washed RBC, makeup to 400 μl with 1x cytomix. After transfection, washed away the broken red blood cells, the mixture was transferred into flasks with medium, and then the parasite schizonts invaded the RBCs that were pre-loaded plasmids, then cultured in the incubator as mentioned above, after 4th days of transfection, BSD and WR99210 drugs were added to the medium until the parasites were killed clearly. Every two days the medium was changed with selected drugs until the parasites reappear and were confirmed by PCR and Sanger Sequencing. Then clones were performed dilution cloning, and the strain that carried the mutant site was further to perform the drug susceptibility.

### Immunofluorescence assay (IFA)

IFA was performed as previously described^30^. Briefly, the asynchronous parasites carrying the 3×HA tag were fixed with 4% paraformaldehyde, then incubated in 1% TrionX-100 at room temperature. Parasites were incubated in PBS buffer containing 5% goat serum (Solarbio, China), then were incubated with mouse anti-HA antibody (Sigma, USA) at 1:500 dilution, after three times with PBS, parasites were incubated with anti-mouse Alexa Fluor 488-conjugated secondary antibody (Cell signaling Technology, USA) at 1:500 dilution and washed with PBS. Then the parasites were incubated with PBS containing 1μg/mL of DAPI. Finally, the images were obtained using a fluorescence microscope.

### *In vitro* drug susceptibility assays

10 antimalarial drugs were used *in vitro* drug susceptibility assays including dihydroartemisinin (DHA), artemether (AM), artesunate (AS), naphthoquine (NQ), melfloquine (MFQ), lumefantrine (LMF), piperaquine (PPQ), naphthoquine (NQ), pyronaridine (PND), chloroquine (CQ) and quinine (QN). *In vitro* drug susceptibility assays were operated following the previous report^31-32^. Briefly, synchronized parasites were adjusted to 0.5% parasitemia and 2% hematocrit at 37°C for 72h with drugs for drug susceptibility assays, then SYBR Green I was used to dye them. The half-maximal inhibitory concentration (IC50) was estimated using a non-linear regression model implemented in GraphPad Prism 6.0.

The ring survival assay of 0-3h (RSA_0-3h_) was also used to evaluate the artemisinin susceptibility of transgenic strains as described previously^33-34^. Briefly, tightly synchronized 0-3 h ring-stage parasites were prepared at 1% parasitemia and 2% hematocrit, then were treated with 700 nM of DHA or the same concentration of solvent (ethanol) for 6 h, then the drug was washed off and cultured another 66 h. Thin blood smears were made to assess the ring survival rates of the strains by microscopy with 10000 RBCs counted on each slide. The ring survival rates were determined by comparing surviving parasites in DHA-treated with those in vehicle-treated wells. Three biological and technical replicates were performed for each parasite isolate. Statistical comparisons were made using the non-parametric Mann-Whitney U test.

### Assessing ACAS activity in *P. falciparum*

The ACAS enzyme-linked immunosorbent assay kit (ELISA) was used to detect the activity of ACAS (MEIMIAN, China). Following the kit instructions, duplicate aliquots of protein samples (pre-adjusted to the same concentration level) and diluent solution were added to the 96-well ELISA plate and incubated at 37°C. Then, the horseradish peroxidase (HRP)-labeled secondary antibody were added and incubated. Subsequently, the chromogenic agent was added, then termination solution was added to each well to stop the reaction, and the readings were taken at a wavelength of 450 nm using a plate reader. Enzyme activity was calculated using the standard curve.

## RESULTS

### PfACAS gene mutant strains including 3D7^Control^, 3D7^S868G^, 3D7^V949I^, 3D7^S868G+V949I^, 129^Control^, 129 ^V949I^, 129 ^S868G+V949I^ were successfully engineered

We successfully engineered PfACAS S868G and V949I variants on 3D7 and the similar background strain 16-129 by CRISPR/Cas9 (Fig. 1). These modified strains including 3D7^Control^, 3D7^S868G^, 3D7^V949I^, 3D7^S868G+V949I^, 129^Control^, 129^V949I^, 129^S868G+V949I^ were successfully achieved gRNAs and PAM motif synonymous mutations (shield-mutation) (Fig. 2) and carried mutations S868G or V949I in both 3D7 and 16-129 strains (Fig. 3).

**Fig. 1.**
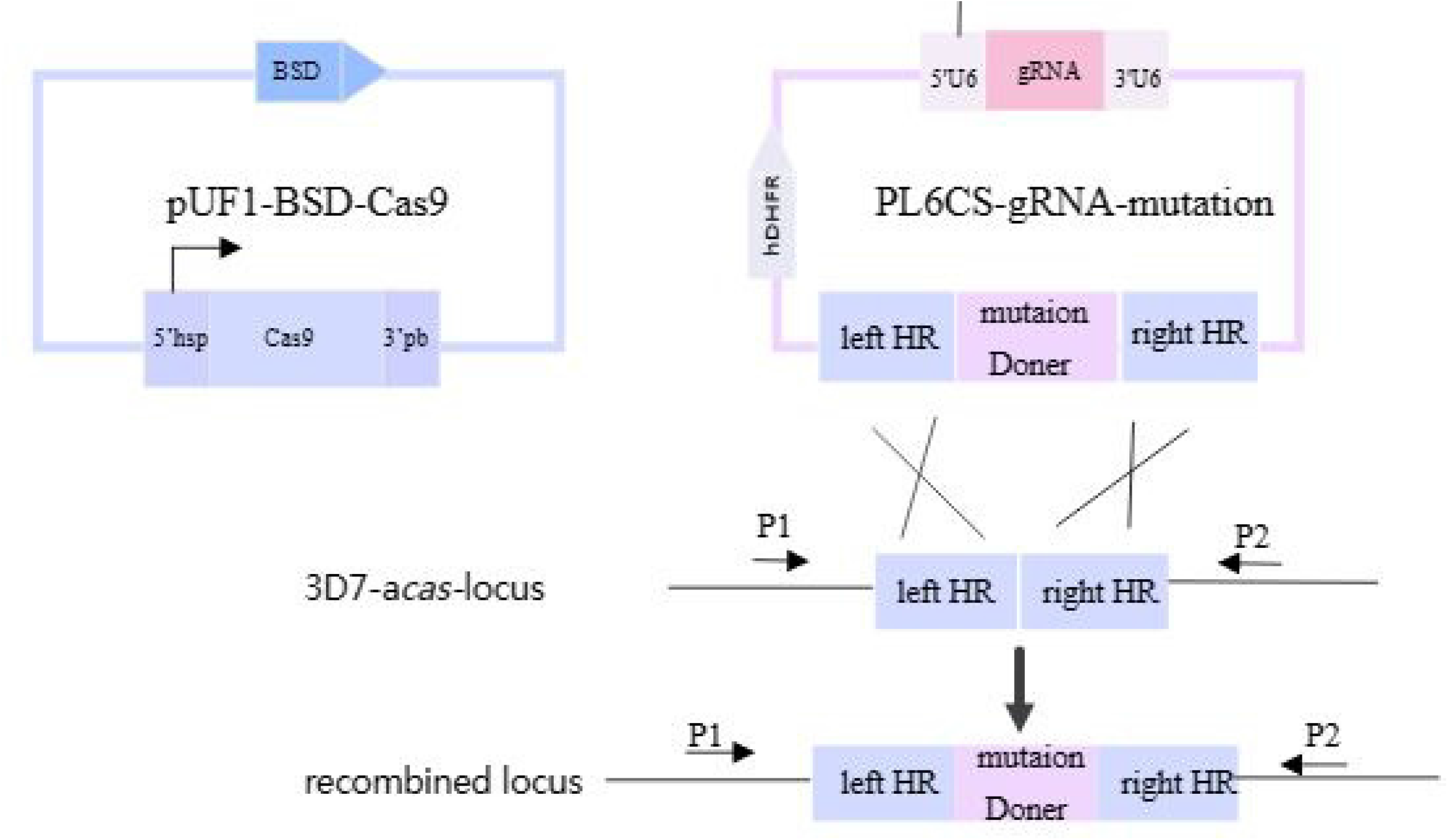
Schematic diagram of labeling the genome using CRISPR/Cas9 gene editing technology. pL6CS-gRNA-mutation plasmid carried mutation Donor and homologous regions (left HR and right HR) and repaired the broken DNA, achieving the introduction of the mutant site into the genome. Primers P1/P2 were used for checking.

**Fig. 2.**
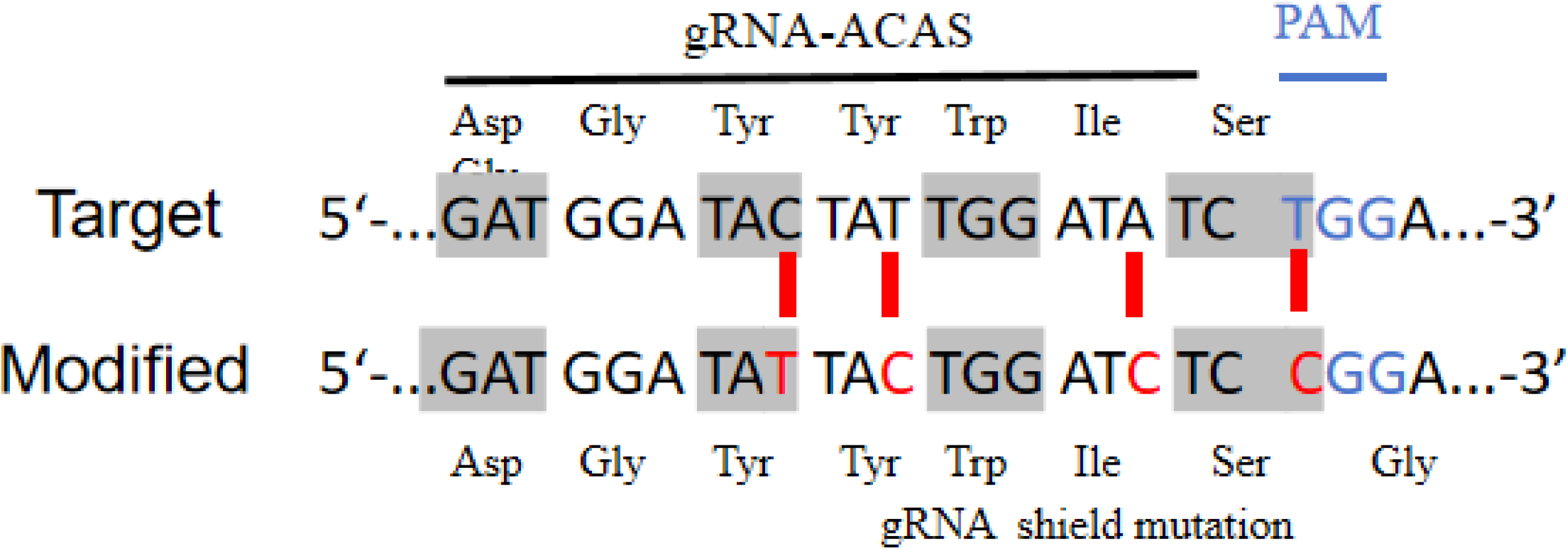
The 20bp length gRNA and PAM (TGG) that were modified as shield-mutations to prevent repeated cutting in the transgenic strains.

**Fig. 3.**
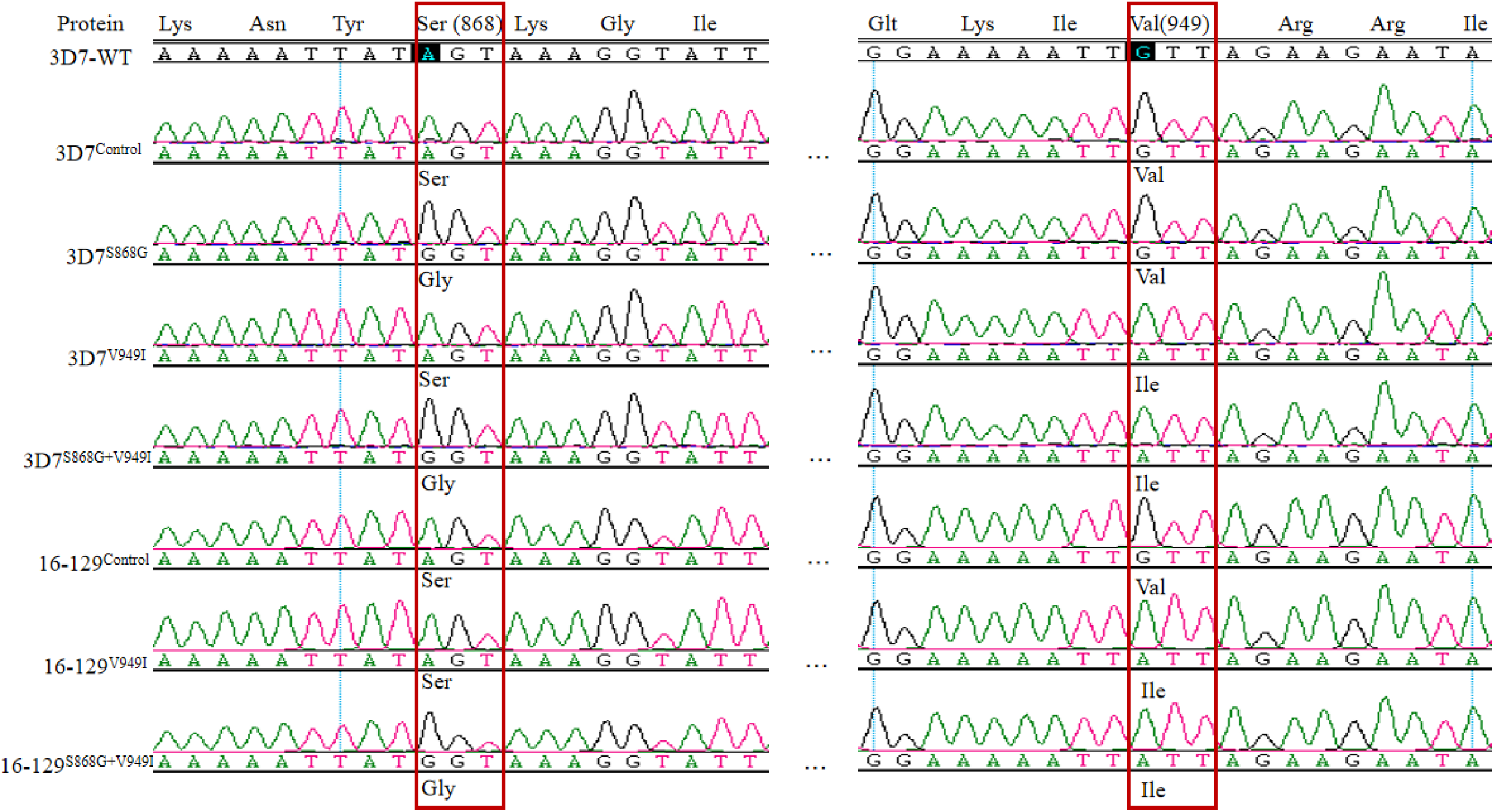
Sequencing in transgenetic strains 3D7 and 16-129, successfully achieved mutations at corresponding positions in the genome of strains.

### *In vitro* susceptibility of engineered strains

*In vitro* susceptibilities to 10 antimalarial drugs of transgenic parasite isolates, the laboratory clone 3D7 and 3D7^C580Y^ and RSA were shown in Table 1. Overall, these engineered lines were sensitive to antimalarial drugs. For CQ, MFQ, QN and PND, the IC50 values of all lineages were lower than the cutoff values that were reported, 100, 30, 600 and 15 nM, respectively ^35^. There were no significant difference between the engineered strains and either 3D7 wild-type or 16-129 wild-type. RSA(0-3h) showed no significant difference between mutant strains and wild type strains, and all strains were lower than 1% (the resistant threshold for ARTR)^33^. For three artemisinin antimalarial drugs, the IC50 values of edited strains were significantly higher than wild-type but showed no significant difference (*P*>0.05). For NQ, PQ and LMF, the IC50 values of engineered lineages were closed to wild-type in 3D7 and 16-129 respectively (Table 1).

**Table 1.**
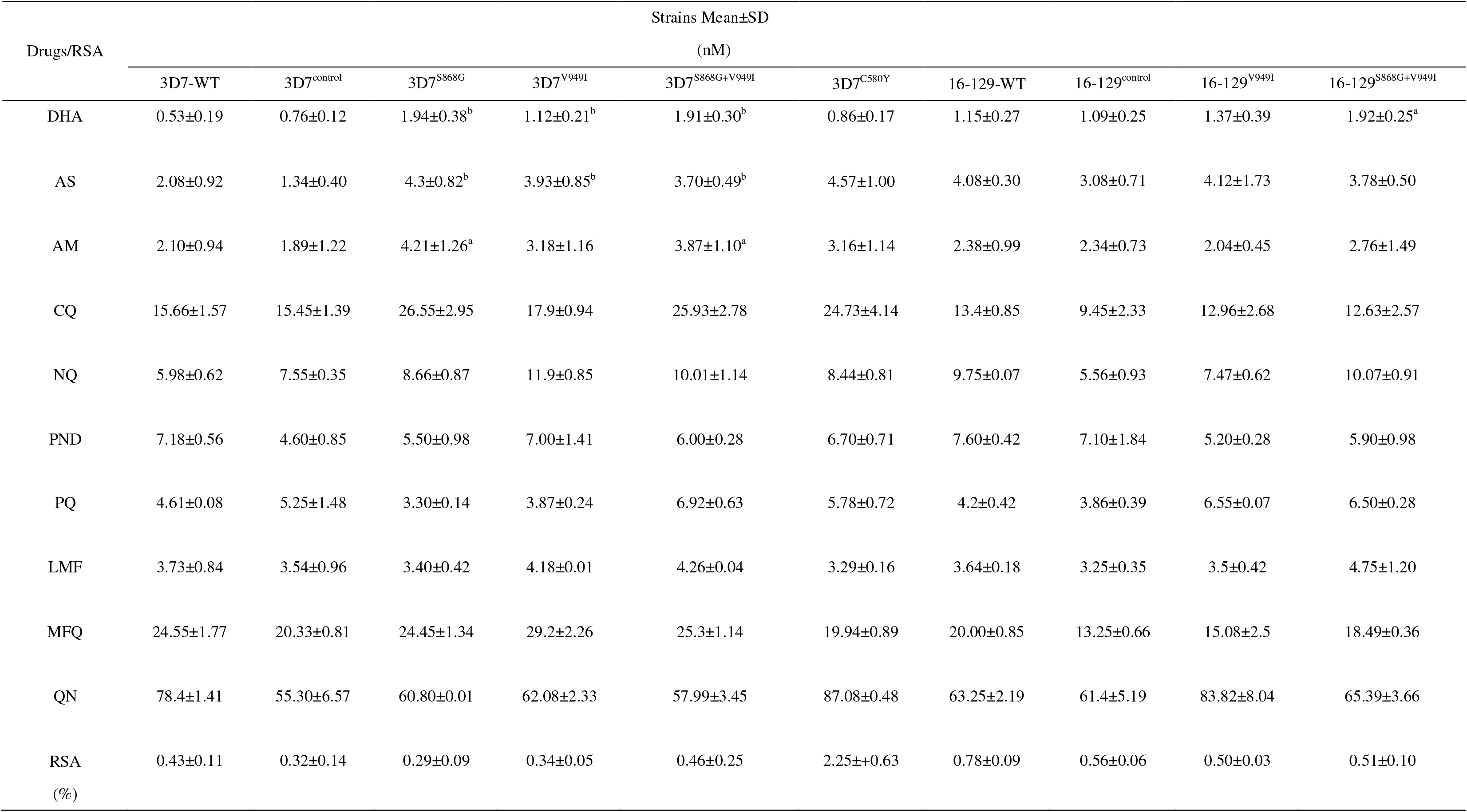
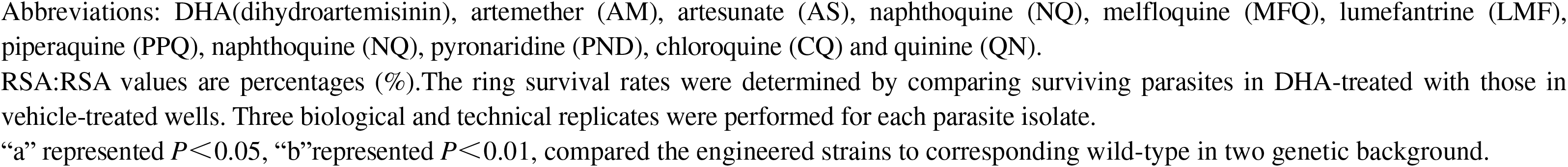
*In vitro* susceptibilities and RSA of studied strains

### ACAS activity and localization of engineered strains

The enzyme-linked immunosorbent assay was used. The ACAS activity of mutant strains results showed no significant difference between the mutant strains and wild type strains (*P*>0.05) (Fig. 4). Through fusion to 3×HA, PfACS could be visualized by fluorescence microscopy, and the HA expression was detected in ring, trophozoite and schizont stages, indicating the presence of PfACAS protein throughout the intraerythrocytic asexual cycle of the parasite, and it was localized in the nuclear and cytoplasm (Fig. S2).

**Fig. 4.**
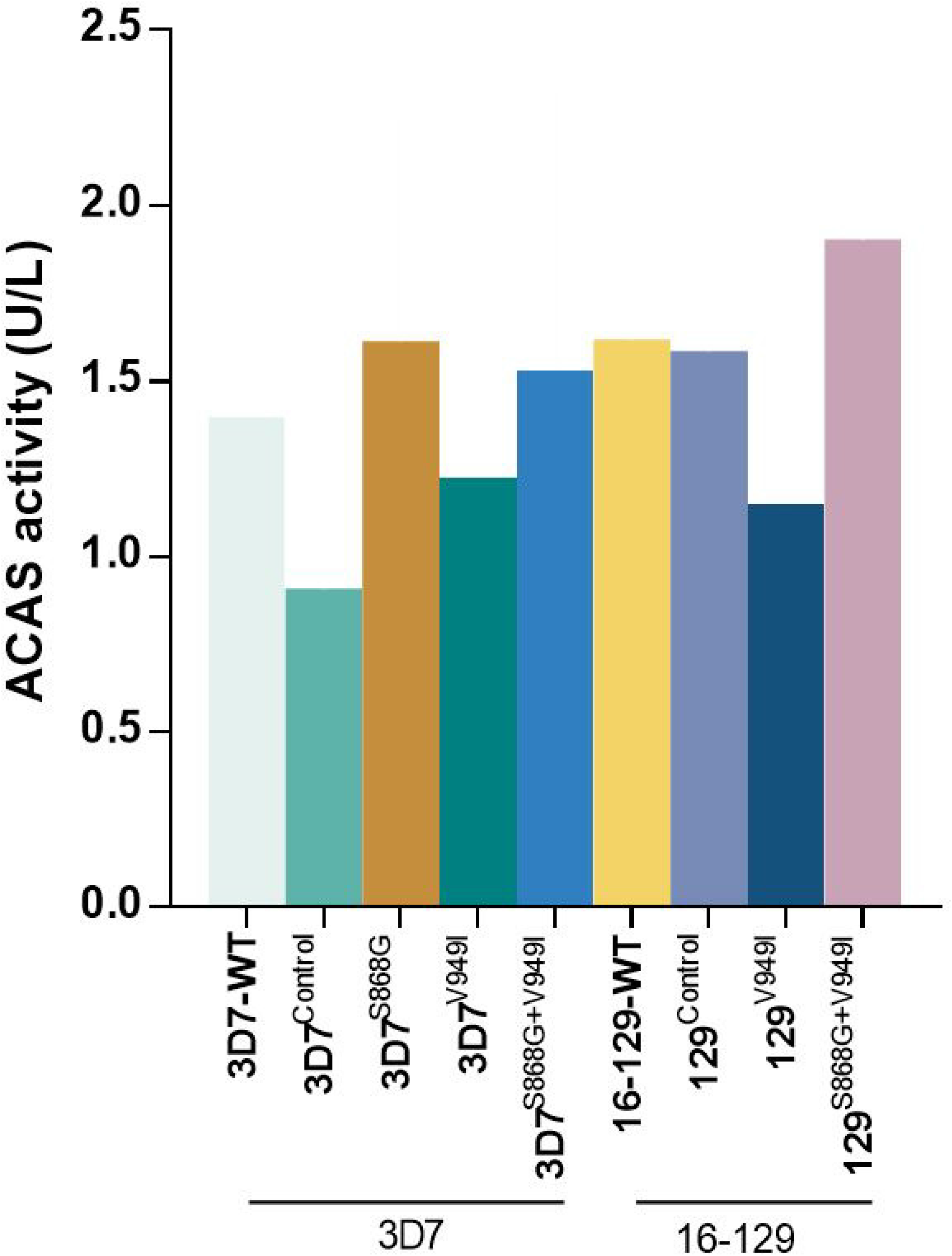
The enzyme activity of 3D7 mutant strains and mutant strains 16-129 of African origin.

## DISCUSSIONS

Here, we wanted to delineate the direct contribution of mutations S868G and V949I in PfACAS gene to antimalarial drugs resistance in *P. falciparum*. We therefore took advantage of a CRISPR/Cas9 gene editing technology and engineered in two genetic background strains, including 3D7 and 16-129. As different strains might behave differently to the same intervention as a result of their distinct genetic background. Furthermore, we evaluated the *in vitro* drug susceptibility assays to 10 antimalarial drugs that commonly used in clinical practice for these engineered strains and examined the ACAS enzyme activity.

Our study shows that these carrying mutations S868G and V949I alone or in combination lines were all sensitive to antimalarial drugs. Summer foundthat there were several mutations in the pfACAS gene (A597V, T648M, Y607C, and A652S),which were edited at DD2 background strain, have been identified that confer the resistance of *P. falciparum* to two small molecules antimalarial compounds (MMV019721 and MMV084978)^25^, whereas they did not detect the reduced susceptibility to several common clinical antimalarials, including DHA, amodiaquine, MFQ. In our study, our mutant strains showed no significant change in sensitivity to 10 antimalarial drugs. This indicates that the resistance characterization of African strains in our study may be due to other factors. Furthermore, the enzyme activity of engineered strains was closed to wild-type, indicating that the mutations did not affect the enzyme’s function. Additionally, immunofluorescence localization experiments showed that PfACAS protein was expressed throughout the entire cycle of *P. falciparum* erythrocytic cycle, which was consistent with the finding of Summers^25^. PfACAS protein was expressed in the nucleus and cytoplasm.

## CONCLUSIONS

The results from this study indicate that mutations in acetyl-CoA synthetase S868G and V949I alone or in combination were insufficient to confer resistance to antimalarial drugs.

## ACKNOWLEDGMENTS

We would like to extend our gratitude to Lubin Jiang team of Key Laboratory of Molecular Virology and Immunology, Chinese Academy of Sciences for kindly providing the pUF1-BSD-Cas9 and pL6CS-Xu-gRNA plasmids and the ARTR strain 3D7^C580Y^.

## AUTHOR CONTRIBUTIONS

Wei Zhao, Methodology, Writing-original draft/ Zhaoqing Yang, Liwang Cui, Conceptulization, Writing-original draft/ Zheng Xiang Methodology, Validation/ Weilin Zeng, Visualization, Data curation/ Yucheng Qin, Maohua Pan, Yanrui Wu, Mengxi Duan,Ye Mou, Tao Liang, Yanmei Zhang, Cheng Liu, Xiuya Tang, Yaming Huang, Gongchao Yang, Investigation, Resources.

## Supplemental Material

Table S1-S3,Fig. S1-2.

## Legends

Fig.S1 A. 3×HA tag sequence (111bp). B. The 3×HA tag sequence was successfully inserted in front of the PfACAS gene stop codon (TAA).

Fig.S2 Immunofluorescence localization of AcAS using an HA Tag. Scale bar: 5μm.

